# Sleeve Gastrectomy enhances glucose utilization and remodels adipose tissue independent of weight loss

**DOI:** 10.1101/779033

**Authors:** David A. Harris, Amir Mina, Dimitrije Cabarkapa, Keyvan Heshmati, Renuka Subramaniam, Alexander S. Banks, Ali Tavakkoli, Eric G. Sheu

## Abstract

**Objective:** Sleeve gastrectomy (SG) induces weight-loss independent improvements in glucose homeostasis by unknown mechanisms. We sought to identify the metabolic adaptations responsible for these improvements.

**Methods:** Non-obese C57Bl6/J mice on standard chow underwent SG or sham surgery. Functional testing and indirect calorimetry were used to capture metabolic phenotypes. Tissue-specific glucose uptake was assessed by 18-FDG PET/CT and RNA sequencing was used for gene expression analysis.

**Results:** In this model, SG induced durable improvements in glucose tolerance despite not causing lasting changes in weight, fat/lean mass, or food intake. Indirect calorimetry revealed post-SG animals had respiratory exchange ratios (RER) nearing 1.0 on average and had daily RER excursions above 1.0, indicating preferential glucose utilization and increased energy demand, respectively. Sham operated mice demonstrate normal RER feeding/fasting excursions. PET/CT showed increased avidity within white adipose depots. Finally, SG led to an upregulation in the transcriptional pathways involved in energy metabolism, adipocyte maturation, and adaptive and innate immune cell chemotaxis and differentiation within the visceral adipose tissue.

**Conclusions:** SG induces a rapid, weight-loss independent shift towards glucose utilization and transcriptional remodeling of metabolic and immune pathways in visceral adipose tissue.

## 1. INTRODUCTION

Bariatric surgery in the form of sleeve gastrectomy (SG) has become the most performed metabolic surgery in the United States. SG leads to rapid and durable improvements in type 2 diabetes (T2D) and this effect often precedes weight loss. [1–4] In fact, nearly 40% of patients with diabetes who undergo SG leave the hospital without needing anti-diabetic medications. [5] Interestingly, there has also been increasing evidence that non-obese, metabolically unhealthy patients can have remission of T2D following surgical intervention. [6] Thus, clinically, there are weight-loss independent, SG-induced adaptations that lead to improved glucose homeostasis.

Multiple pre-clinical and clinical studies aimed at identifying the molecular basis for post-SG T2D remission show that surgery leads to complex changes in incretin hormone and bile acid production, changes in gut microbial composition, intestinal cellular adaptation, and immune modulation. [7–9] However, multiple authors have revealed that while these processes play a role in post-surgical T2D remission, they are not sufficient, in-and-of-themselves, to cause remission. [10] [11–13]

Furthermore, there has been little directed effort to identifying which of these changes, if any, are responsible for the early, weight-loss independent T2D improvement seen clinically. Investigators have tried to address this by using sham pair-feeding and have shown that SG can lead to adipose adaptive immune cell changes, improved muscle and hepatic insulin sensitivity, and reduced hepatic steatosis in the absence of significant weight loss. [9,14,15] However, one major limitation to these studies is that they are derived in obese models where animals lose weight following surgery and are maintained on non-physiologic, high-fat diets indefinitely. This makes it increasingly difficult to define the SG-specific, weight-loss specific, and diet-specific changes that are occurring after SG. Thus, currently the weight and diet independent metabolic adaptations and target tissue of SG remain poorly defined.

Here, we describe a novel model of SG in non-obese mice and show that SG leads to weight independent changes in glucose tolerance. This post-SG phenotype is defined by a global metabolic shift towards increased glucose utilization and appears to lead to, or be the result of, visceral adipose tissue (VAT) immunologic remodeling, glucose sequestration, and glucose use.

## 2. MATERIALS AND METHODS

### 2.1 Animals

11-week-old, male, C57Bl/6J mice were purchased from Jackson Laboratory (Bar Harbor, ME). They were housed in a climate-controlled environment with a 12-hour light/dark cycle and reared on a normal rodent chow (5% calories from fat; Pico5053, Lab diet, St. Louis, MO). They acclimated for 1 week prior to undergoing any procedure. Figure 1 outlines the overall experimental design. All procedures were approved by the Institutional Animal Care and Use Committee and animals were cared for according to guidelines set forth by the American Association for Laboratory Animal Science.

**Figure 1.**
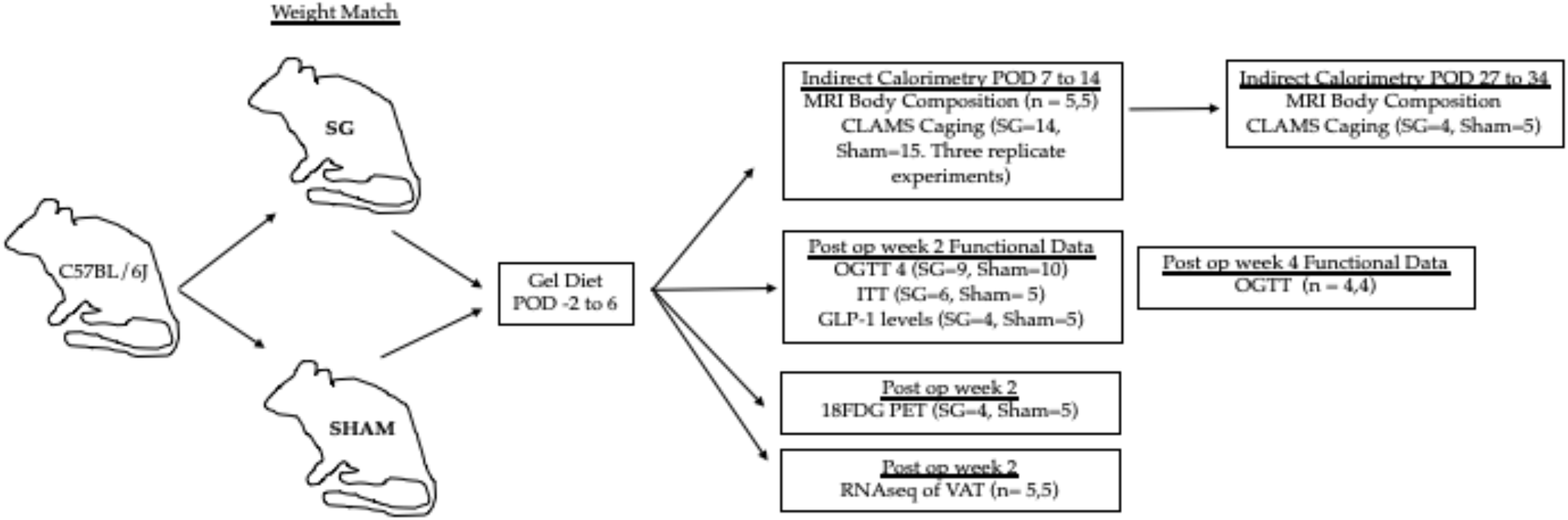
Experimental design.

### 2.2 Sleeve gastrectomy (SG) and sham procedures

Mice were weight-matched into two groups and either underwent SG or Sham operation. Under isoflurane anesthesia, SG and Sham procedures were performed via a small midline laparotomy. In SG, the stomach was dissected free from its surrounding attachments, the short gastric vessels between the stomach and spleen were divided, and a tubular stomach was created by removing 80% and 100% of the glandular and non-glandular stomach, respectively. Sham operation consisted a similar gastric dissection, short gastric vessel ligation, and manipulation of the stomach along the staple line equivalent. Short gastric vessel ligation was chosen as part of our control as ligation of these vessels effect portal venous flow. Signaling through the portal vein has been shown to be important in the body’s ability to sense and respond to glucose. [16,17] Mice were then individually housed thereafter to allow for monitoring of food intake, weight, and behavior. After surgery, SG and Sham mice were maintained on Recovery Gel Diet (Clear H_2_O, Westbrook, ME) for 6 days and then restarted on normal chow.

### 2.3 Functional glucose testing

Oral glucose tolerance testing (OGTT) was performed on post-operative week 2 and 4 and insulin tolerance testing (ITT) was performed on post-operative week 2. Animals underwent a 4 hour fast (8 am to noon). During OGTT, mice received 2mg/g of oral D-Glucose (Sigma-Aldrich, St. Louis, MO) and serum glucose levels were measured from the tail vein at 15, 30, 60, and 120 min with a OneTouch Glucometer (Life technologies, San Diego, CA). ITT was performed by intraperitoneal instillation of 0.6U/kg of regular human insulin (Eli Lily and Company, Indianapolis, IN) and measurement of serum glucose levels at 15, 30, and 60 min. Baseline glucose was measured for each animal prior to medication administration. An ELISA assay was used to quantify the concentration of total glucagon like peptide-1 (GLP-1) in circulation 15 minutes following glucose gavage as above (Crystal Chem, Cat81508, Elk Grove Village, IL). Each functional assay represents a separate animal experiment.

### 2.4 Body composition analysis and Comprehensive Animal Monitoring Systems (CLAMS)

A separate group of SG and Sham mice were placed into CLAMS systems from post-operative day 7 to 14 and then again on post-operative day 27 to 34. Body composition of each mouse was determined by MRI spectroscopy. Mice were then placed into individual temperature-controlled cages within the CLAMS system, which was maintained with a normal 12-hour light/dark cycle at 22°C. Mice had access to food and water ad libitum. Individual mouse CO_2_ expenditure, O_2_ consumption, food consumption, and locomotor and ambulatory were collected over the week. The first 24 hours of each experiment was discarded to account for mouse acclimation. Energy expenditure and the respiratory exchange ratio were calculated. CLAMS data was analyzed using the open source program CalR as previously described [18]. Experimentation from post-operative day 7 to 14 was completed in triplicate and the data was pooled for analysis (Sham n =14; SG n= 15). A single longitudinal study was completed for CLAMS caging (post-operative day 27 to 34, sham n = 5, SG n = 4).

### 2.5 PET/CT imaging with 18-Fluoro-deoxyglucose

Two separate experiments were completed to explore glucose handling in SG mice. Two weeks following SG (n=2) or sham (n=2) operations, mice underwent PET/CT imaging with orally delivered 18-FDG. Each mouse was administered 200μl of 18-FDG such that there was equivalency in the dose of molar glucose given. Under isoflurane anesthesia, PET/CT experimentation was completed over multiple days so that the activity of delivered 18-FDG varied between 116 and 300μCI. Using PMOD software (PMOD technologies LLC, Zurich, Switzerland) time activity curves were created over a 60 minutes and PET imaging was overlaid onto captured CT images. In a second replicate experiment SG (n=2) and sham mice (n=3) were treated similarly. Mice were euthanized and whole blood and organs were placed into a well counter to assess for overall activity. Data from well counting are pooled between the two experiments (SG n=4 and sham n=5). Comparisons between organ SUV was completed in GraphPad Prism.

### 2.6 RNAseq of visceral fat

Two weeks following SG (n=5) or sham (n=5) operations, mice were euthanized and visceral fat (intra-abdominal, epididymal fat pad) was collected. [19] Total RNA was extracted according to the manufacturer’s instructions (Qiagen, RNeasy Lipid Tissue Mini). RNA quality was ensured using an Agilent Bioanalyzer (Agilent Technologies, Santa Clara, CA). Libraries were prepared using Roche Kapa RiboErase rRNA depletion stranded totalRNA Hyper Prep sample preparation kits from 100ng of purified total RNA according to the manufacturer’s protocol. The finished dsDNA libraries were quantified by Qubit fluorometer, Agilent TapeStation 2200, and RT-qPCR using the Roche Kapa library quantification kit according to manufacturer’s protocols. Uniquely indexed libraries were pooled in equimolar ratios and sequenced on an Illumina NextSeq500 with paired-end 75bp reads by the Dana-Farber Cancer Institute Molecular Biology Core Facilities. Sequenced reads were aligned to the UCSC mm9 reference genome assembly and gene counts were quantified using STAR (v2.5.1b). [20] Differential gene expression testing was performed by DESeq2 (v1.10.1) and normalized read counts (FPKM) were calculated using cufflinks (v2.2.1). [21,22] RNAseq analysis was performed using the VIPER snakemake pipeline. [23]

The EnrichR platform was used to assess gene expression ontology. [24,25] Functional enrichment with WikiPathways 2019 Mouse and ChEA2016 Transcription pathway analysis are reported.

### 2.7 Statistics

GraphPad Prism 8 (San Diego, Ca), CalR statistical software [18], and the Enrichr platform (http://amp.pharm.mssm.edu/Enrichr/) were used for data analysis. Student’s t-tests were used for continuous variables. General linear modeling was used were appropriate with a lean mass covariate. Statistical significance is denoted in figures and text. For RNA-seq, gene ontology and transcriptional pathways, adjusted p-values<0.05 are listed in order of Enrichr combined score, which is the product of the log of the Fischer’s exact p-value and the z-score (deviation from expected rank).

## 3 RESULTS

### 3.1 Weight-loss independent improvements in glucose metabolism following sleeve gastrectomy

To investigate the weight-loss independent, anti-diabetic effects of SG, we developed a novel model of SG in 11-week old C57Bl/6J mice reared on standard chow. After a 1-week acclimation period, mice underwent either SG or sham operation. While there was an initial difference in weight loss in SG mice as compared to sham control animals, by post-operative day (POD) 7, mice from both groups showed identical weights (Figure 2A). Mice were restarted on solid chow on POD 6. By POD 7, overall caloric intake as measured by daily food weight was not different between groups (Figure 2B). Further, MRI body composition analysis revealed that SG mice had reduced fat and lean mass during the first operative week, but by POD 14, all mice had identical body composition (Figure 2C). Thus, this model is well suited to study the weight-loss independent effects of SG.

**Figure 2:**
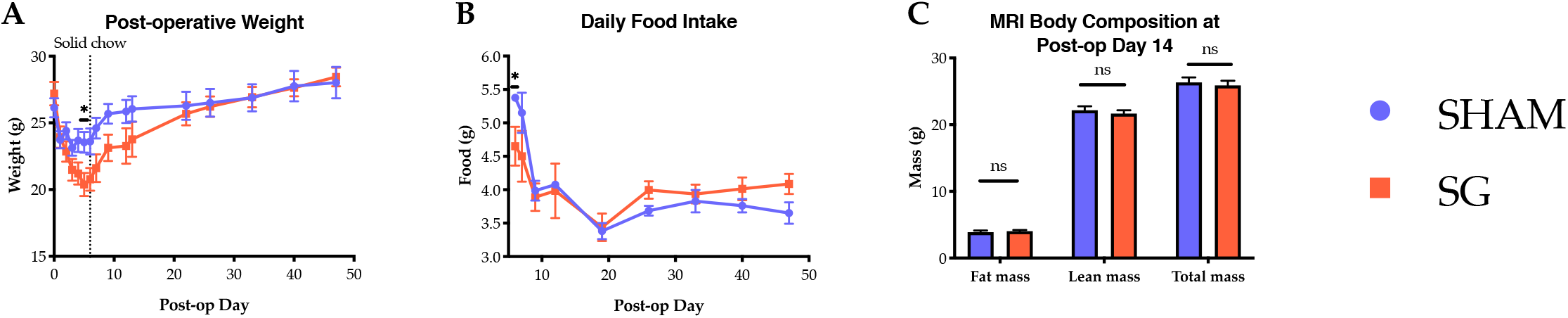
Post-operative weight (A), daily food intake (B), and body composition (C) between Sham (n=4) and SG (n=4) mice. A and B, n = 4,4. For C, n=5,5 and data is representative of 3 biologic replicates. Comparisons by Students’ t-tests. *p<0.05

Despite equivalency in weight, caloric intake, and body composition by post-operative day 14, SG mice displayed distinctly different glucose handling compared to their Sham counterparts. Oral glucose tolerance testing was performed in the second post-operative week. While SG and Sham animals had no difference in fasting or maximal glucose peak, glucose levels in SG animals rapidly cleared, with serum levels dropping nearly 200 mg/dL to baseline levels in the 15 min following maximal glycemia. This change was still evident at 4 weeks (Figure 3A and B). Additionally, SG animals had a more rapid decline in serum glucose levels during ITT 2 weeks post-surgery; although, there was no difference in AUC between groups (AUC Sham vs. SG, 5527 ± 851 vs 3792 ± 598, p = 0.127). Of note, during this testing, two SG animals and one Sham had to be rescued from hypoglycemia at 60 minutes. In keeping with the known post-SG physiology in humans, SG mice had increased serum GLP-1 levels in response to oral glucose stimulation at 15 minutes (Figure 3d). These data demonstrate improved glucose homeostasis in mice following SG, independent of differences in body composition.

**Figure 3:**
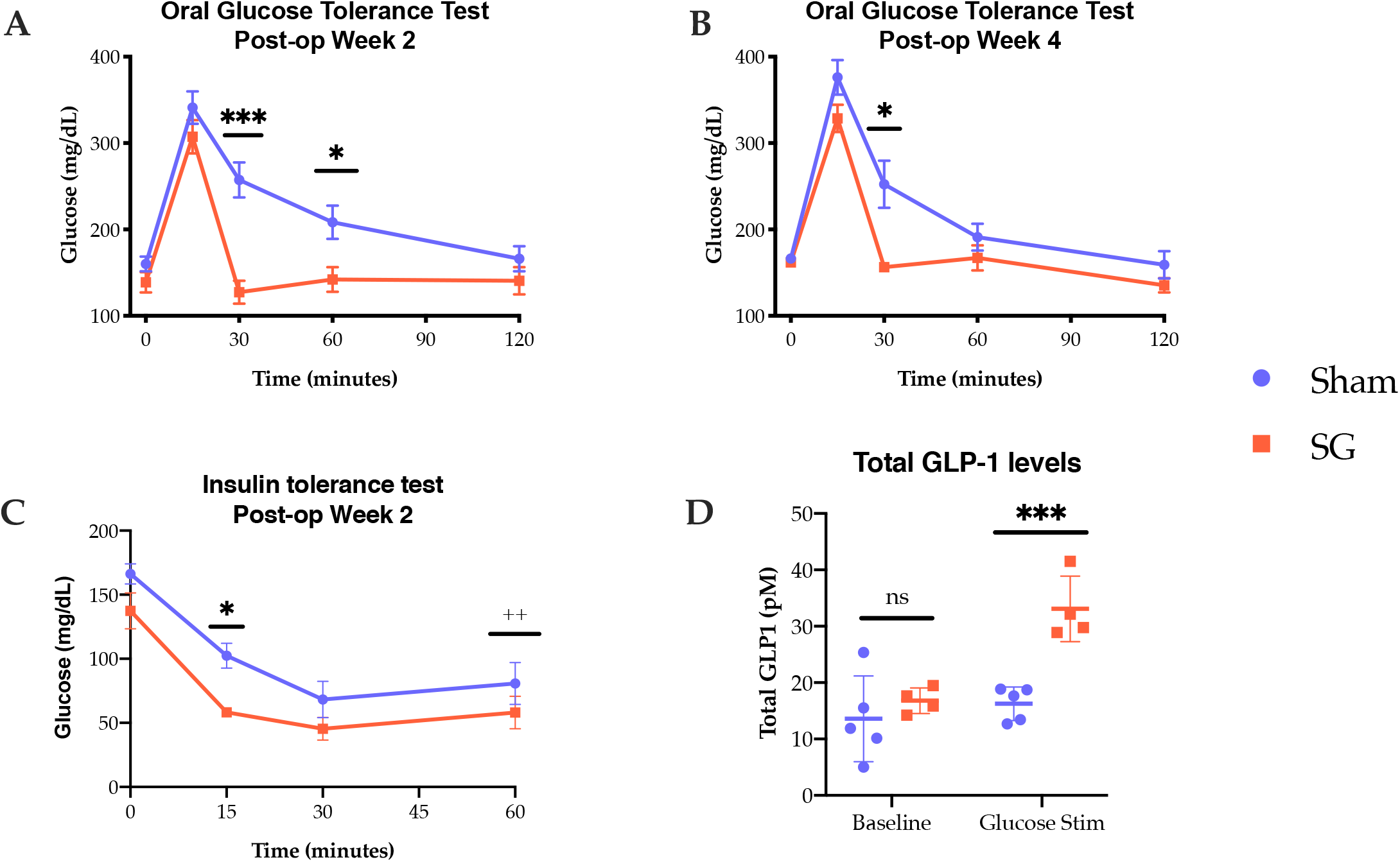
(A,B) Oral glucose tolerance testing at 2 and 4 weeks (A; SG n=9, Sham n=10, B; n=4,4). (C) Insulin tolerance testing at 2 weeks (SG n=6, Sham n=6). (D) Total GLP1 levels at 15 minutes following glucose challenge (SG n=4 and Sham n=5). Comparisons made using t-tests. *p<0.05, **p<0.01,***P<0.001. ++ One sham and two SG animals had to be rescued from hypoglycemia at this time point.

### 3.2 Indirect calorimetry reveals increased carbohydrate utilization in SG mice

To gain a better understanding of the metabolic changes underlying the observed weight-loss independent improvement in glucose handling by SG mice, we next utilized indirect calorimetry via Comprehensive Lab Animal Monitoring Systems (CLAMS) to capture the energy balance of these animals. SG and Sham mice were placed into CLAMS cages from POD 7 to 14. The most notable change occurred in the respiratory exchange ratio (RER; Table A1). As typical of mice fed a standard rodent chow, the RER of Sham animals ranged from 0.85 to 1.0 during light and dark photoperiods corresponding, roughly, to periods of fasting and feeding, respectively. In contrast, SG mice had a strikingly high RER ranging from an average of 0.9 to an average peak near 1.05. RER values greater than 1.0 suggest either increased glucose utilization or a process such as de-novo lipogenesis (Figure A and B). [26] Importantly, there was equivalency in the total food consumed in a 24-hour period and ambulatory activity (Table A2). During the first post-operative week, SG mice showed a reduction in the EE across all times of day (Figure 4C, Figure A3). The effect on EE reduction is consistent with effects seen in food restriction experiments for weight loss in rodents and humans. [27,28] However, the effect on RER is novel.

**Figure 4:**
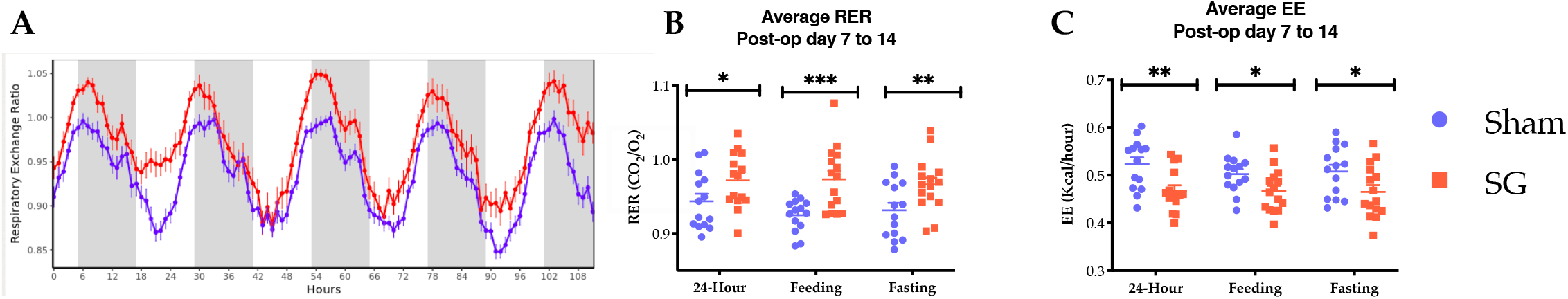
SG mice have continuously elevated RERs (average 0.9 to 1.05) indicating preferential glucose utilization during both feeding and fasting. Sham mice demonstrate normal RER excursions, reflective of mixed lipid/glucose utilization (A). SG mice have higher average RER (B) and lower EE over a 24 hour period, while feeding, and while fasting (C); Combination of 3 biologic replicates. SG n =15 and sham n=14; t-test, *p<0.05, **p<0.01, ***p<0.001

After two additional weeks of normal housing and daily monitoring, mice were again placed into CLAMS cages (POD 27 to 34). At this point, SG and Sham mice had similar total (26.3±0.31 vs. 25.4±0.33, p=0.58), fat (4.1±0.3 vs. 4.0±0.1, p=0.70), and lean masses (22.0±0.9 vs 21.6±0.3, p=0.53). During this week, SG and Sham mice had the same caloric intake, the same overall energy expenditure, and same activity level. However, the elevation in the RER after SG persisted (Table A4) indicating that changes in RER are durable and important in the underlying physiology of SG.

### 3.3 Positron-emission tomography with oral 18-FDG reveals a tissue-glucose sink

We next sought to identify the tissue responsible for rapid systemic glucose utilization during OGTT and for the RER changes seen in SG mice. As show in Figure 5a, oral delivery of 18-FDG to SG mice led to rapid distal delivery of the glucose analog when compared to Sham animals. Time activity curves generated from PET/CT imaging revealed that the glucose analog remained stagnant in the stomach and very proximal small bowel in Sham animals, while in SG animals there was rapid gastric clearance and delivery to the distal small bowel within 60 minutes (Figure 5B).

**Figure 5:**
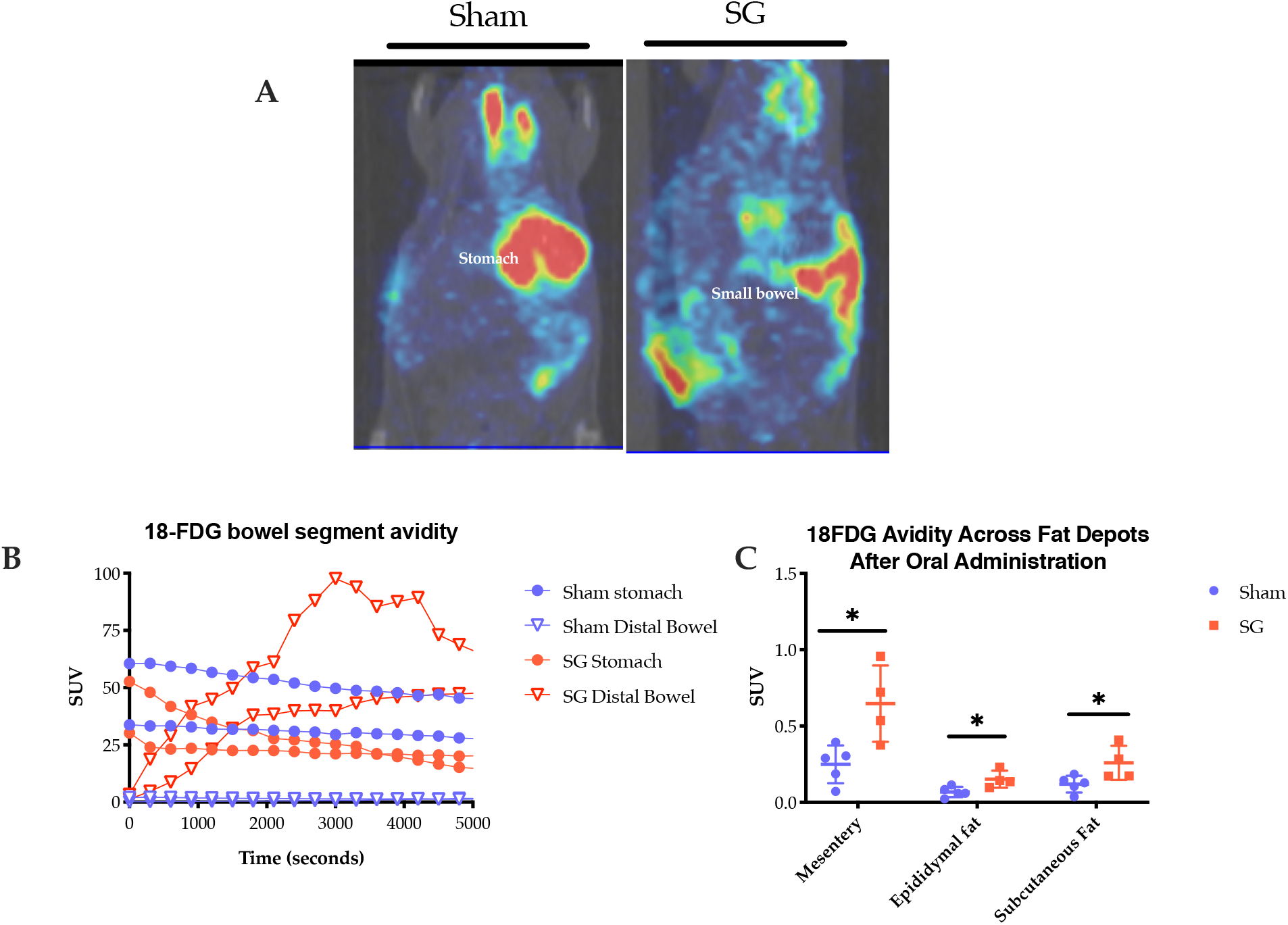
(A) Representative PET/CT images from Sham (left) and SG (right) animals 1 hour after oral 18-FDG administration during post operative week 2. Red denotes high and green, low 18-FDG avidity. (B) Time activity curves were generated (n=2,2). (C) 18-FDG avidity across all fat depots as measured by well counter (C: SG n=4, sham n=5). *p<0.05

Quantification of 18-FDG avidity by well counter at 90 minutes showed that most tissues had equivalency in total glucose uptake (Table A5). There were no statistically significant differences in bowel segment avidity. However, there was a significant increase in the avidity of cecal stool in SG animals suggesting that there is increased luminal transit, as opposed to an increased enterocyte uptake and capture of 18-FDG. However, in SG animals, there was increased 18-FDG present across all fat depots tested – mesenteric, epididymal, and subcutaneous – suggesting that metabolic changes within these fat depots could explain the post-SG physiology.

### 3.4 RNA sequencing reveals gene level changes in visceral fat immunity and metabolism

Metabolic dysregulation and inflammation of the visceral adipose tissue (VAT) are known, potent contributors to the pathogenesis of metabolic syndrome. Thus, changes in gene expression and transcriptional programming in this depot may underlie the post-SG improvements. Given the increase in 18-FDG in the VAT of SG mice and its known importance to metabolic regulation, we utilized RNA sequencing to better understand the gene expression profile and transcriptional networking of this depot in SG.

Table 1 outlines significant gene-level changes following SG. There was a large shift in the immunologic gene expression profile of post-SG VAT with increased expression of CXCL13 and CCL8, CISH, and IgJ corresponding to up-regulation of B and T cell chemotaxis, regulation of T cell activation, and immunoglobulin cross linking, respectively. We also observed changes in the VAT metabolic gene profile with an upregulation of EHHADH – a peroxisomal enzyme responsible for beta oxidation of fatty acids – and related processes of fatty acid beta-oxidation, peroxisome function, and amino acid metabolism. In keeping with the published human data showing decreased systemic and organ-specific leptin levels following sleeve gastrectomy, there was a reduction in the leptin (LEP) expression within the VAT of SG animals [29–31]. There was also a reduction in oxytocin receptor signaling, which partially controls adipogenesis and fat accumulation in fat-tissue depots [32].

**Table 1.** Top changes in relative gene expression in the visceral fat of SG compared to Sham animals. Base >100, Fold change >1.0, and p<0.05. n=5,5

| <u>UPREGULATION</u> |  | <u>DOWN REGULATION</u> |  |
| --- | --- | --- | --- |
| Ehhadh | Ldb3 | Ehd3 | Nnat |
| Cxcl13 | Cish | Oxtr | Mroh2a |
| Igj | Ccl8 | Lep | Sfrp5 |
|  |  | Atp6v0d2 |  |

Next, pathway analysis was completed with the use of WikiPathways 2019. Again, there was an upregulation of immune-centric pathways including phagocytosis, IL-2, IL-3, IL-5, chemokine signaling, and TYRO protein tyrosine kinase binding protein signaling. The latter is associated with immune cell maturation and activation (Table 2). There were no pathways that were down regulated.

**Table 2.** Upregulated specific pathway changes identified in SG animals through functional enrichment analysis using WikiPathways 2019 Mouse. n=5,5

| <u>PATHWAY</u> | <u>P-VALUE</u> | <u>COMBINED SCORE</u> |
| --- | --- | --- |
| Chemokine Signaling | 0.023 | 76.7 |
| TYROBP Causal Network | 0.024 | 62.0 |
| Microglia Pathogen Phagocytosis | 0.010 | 56.0 |
| Macrophage Markers | 0.036 | 45.6 |
| IL-2 Signaling | 0.001 | 45.1 |
| IL-3 Signaling | 0.001 | 29.2 |
| IL-5 Signaling | 0.036 | 15.7 |

Finally, transcription factor binding sites upstream of theses target gene sets were identified via ChEA2016 pathway analysis (Figure 6). The gene expression profile in SG VAT mapped to traditional master-regulators involved in lipid and glucose homeostasis – peroxisome proliferator-activated receptor gamma (PPARγ) and forkhead box protein O1 (FOXO1) –responsible for diverse processes including glucose metabolism and adipocyte cell fate. There was contribution from silencing mediator of retinoid and thyroid hormone receptors (SMRT) and nuclear receptor co-repressor (NCOR), which through subdomains, effect PPARy signaling and modulate the immune response by transrepression of co-activators of NF-kB, IRFs and LPS targets. [33–36] Finally, there was expression mapping to other transcription factors such as Interferon-regulatory factor 8 (IRF8), MDS1 and EVI1 Complex Locus (MECOM), Polycomb Repressive Complex 2 Subunit (SUZ12), and E2A immunoglobulin enhancer-binding factor E12/E47 (E2A), which have roles in determining immune cell fate, cell resource utilization, and/or other metabolic processes.

**Figure 6:**
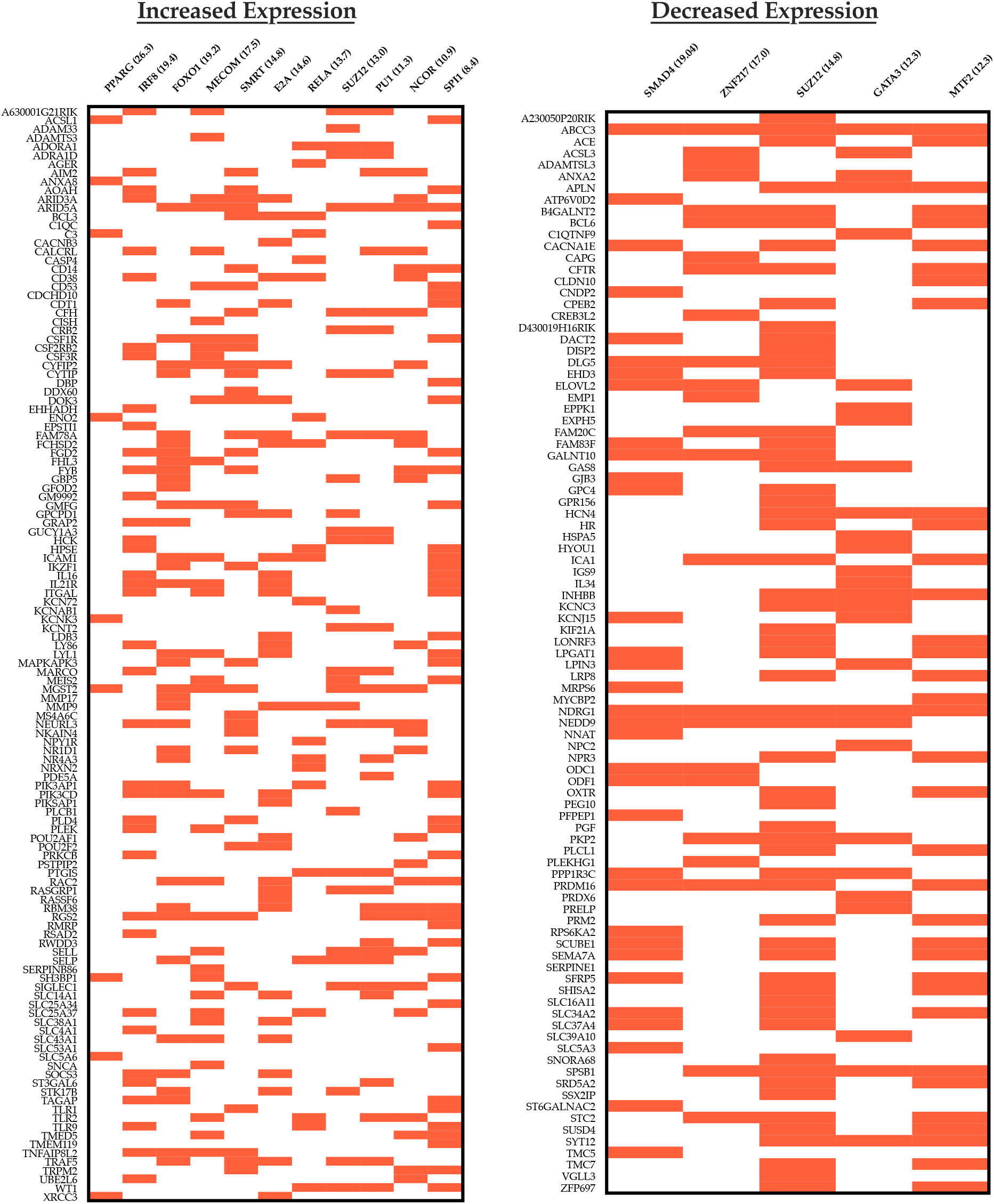
ChEA2016 transcription pathway analysis showing the most up regulated (left) and down regulated (right) transcription pathways. Data is organized as clustergrams with a red bar denoting a gene contributing to the pathway expression. Only pathways with an adjusted p<0.05 are included. Pathway titles are followed by the combined score. Base > 50, log change > 0.5, p<0.05

Transcriptional program analysis of downregulated pathways revealed a shared SUZ12 signature, suggesting that alternative expression of downstream products may be important to the global change in fat homeostasis and inflammation following SG. This is further supported by downregulation in the Metal Response Element Binding Transcription Factor 2 (MTF2), which guides SUZ12 to specific promoters. [37] Finally, there was a decrease in gata binding protein 3 (GATA3) which is classically associated with a TH2 immune response but is also prevalent in adipocyte precursors. GATA3 also suppresses PPARy, so its downregulation is in keeping with the observed increase PPARy expression identified in the VAT of SG mice. [38]

## 4 DISCUSSION

The immediate benefits of SG on glucose metabolism and insulin resistance has been documented in both human patients and mice. We have previously shown that within days of surgery, roughly 40 percent of human patients have durable improvement, or resolution, in their diabetes. [5] The mechanisms behind these early, rapid changes are unknown but are critical to understanding the benefits of surgery. Our study in mice suggests that SG leads to an increase in visceral fat glucose uptake and global glucose utilization, which either drives, or is the sequela of, immunologic remodeling. We postulate that these processes likely underlie immediate post-SG improvements in glucose handling.

To date, most murine studies of SG have utilized a diet-induced model of obesity (DIO) or a genetically obese model, combined with sham pair-feeding, in an attempt to recapitulate and study SG physiology. [9,11,14,15,39–41] While mice in these studies become obese and have hyperglycemia similar to the target patient population, investigations are limited by differences in food intake via pair-feeding and/or differences in body weight between the surgical intervention and sham groups. Furthermore, in most experiments, pair feeding induces weight loss. Here, we show that the immediate, weight-loss independent effects of SG can be modeled in lean, wildtype, C57Bl/6J mice reared on standard rodent chow. In this model, SG and sham mice have similar food intake profiles. Thus, this model decouples the effects of surgery from changes in diet and body composition.

In our model, SG and sham animals showed equivalent weight, body composition, and daily food intake without pair feeding. Despite this, SG animals had improvements in glucose homeostasis, insulin resistance, and increased circulating GLP-1; all of which are hallmarks of post-SG physiology. [30] Specifically, when challenged with oral glucose, SG animals showed a dramatic reduction in systemic glucose levels 15 to 30 minutes post gavage with a corollary increase in serum GLP-1 expression at 15 minutes. This “glucose sink” phenomenon suggested increased tissue glucose uptake and utilization. In addition, this adds to the growing literature that SG, in the absence of weight loss, leads to changes in metabolic physiology. [9,14,15]

Indirect calorimetry corroborated these findings. The RER, or the ratio of CO_2_ produced to O_2_ consumed, is a function of the substrate of cellular respiration. When an animal is predominantly using carbohydrates as a fuel source, the RER approximates 1.0 as every molecule of O_2_ consumed produces a single molecule of CO_2_. When an animal uses fatty acids as a source of respiration, the RER approaches 0.7. During a typical 24-hour cycle, sham mice vacillated between 0.85 and 1.0 corresponding to fasting or feeding, respectively. However, SG animals had a durable increase in the RER which neared 1.0 during both night and dark cycles indicating preferential glucose utilization during both fasting and feeding. Thus, in SG mice there appears to be an early cellular adaptation to increased glucose allocation and utilization likely leading to long term improvements as shown by the continually elevated RERs in SG mice 4 weeks following surgery.

Interestingly, calorimetry also revealed that SG mice had daily RER excursions above 1.0 and nearing 1.1. This phenomenon has been described previously in maximally exercising individuals as a surrogate marker for attainment of the maximum VO_2_ and in metabolically expensive processes, such as de novo lipogenesis (DNL), owing to its reliance on the pentose phosphate pathway to produce reducing equivalents. [42] [26,43] Studies of RER in humans and mice following SG are limited. Dereppe et al., have shown that human patients following bariatric surgery have an increase in the maximal RER attained (up to an average of 1.28) while exercising. [44] However, SG mice in the current study, do not show an increase in ambulatory activity to suggest exercise, or the equivalent, as the main driver of RER elevations.

PET-CT imaging showed increased 18-FDG uptake across all white adipose depots in SG compared to sham animals suggesting that these depots are, in part, responsible for the glucose sink phenomenon and the elevations in RER. Importantly, this change occurred in the absence of changes in overall, fat, and lean mass and thus appear surgery specific. In contrast to Saedi and colleagues who found that intravenously delivered 18-FDG was sequestered to the alimentary roux-limb of rats following Roux-en-Y gastric bypass, we found no difference in bowel specific glucose uptake in SG animals. [45] These differences may be related to the route of 18-FDG administration. However, owing to the importance of incretin hormones in post-SG outcomes, we felt that orally administered glucose was more physiologically relevant. These differences highlight that while these two procedures may result in clinically and physiologically similar outcomes, they may differ in how those outcomes are achieved.

Based on these findings we explored VAT gene-level changes in SG mice. This revealed an upregulation in multiple metabolic processes that potentially indicate increased lipolysis, increased de novo lipogenesis, and modulated adipocyte cell fate. For example, upregulation in FOXO1 pathways and the gene EHHADH indicates increased lipolysis and beta oxidation of long chain fatty acids, respectively. [46,47] SG animals also had an upregulation in MECOM regulated pathways, which are associated with increased purine and pyrimidine metabolism, amino acid metabolism, pentose phosphate pathway reliance, and glycolysis. [48]. Elevations in PPARγ, SMRT, and NCOR which all interact to control downstream adipocyte fate and function, glucose handling, insulin sensitivity, mitochondrial oxidative capacity, and thermogenesis, were also found. [43,49] [36,50] [33] [51–53]. Thus, as inferred from the calorimetry data, there is upregulation of transcriptional machinery capable of increasing energy production and glucose utilization within the VAT of SG mice.

Unexpectedly, SG mice had an upregulation in multiple immune processes within the VAT. Most of the isolated, gene-level changes occurred in pathways for myeloid and B cell chemotaxis and differentiation. Increased levels of CXCL13 and IgJ suggests B cell chemotaxis to SG VAT and increased immunoglobulin production and crosslinking. Upregulation of transcription factors outlined in Figure 6 are likely underlying these changes. [54] These again include NCOR and SMRT, which both coordinate to modulate the immune response through NF-kB, IRFs and LPS target genes and reduce the pro-inflammatory phenotype of macrophages. [34,35]. We also identified a robust signal in IRF8, which is associated with innate and adaptive immune response and adipocyte metabolic regulation [55] [56,57] and SUZ12, which has been shown to be a regulator of energy metabolism through modulation of brown fat thermogenesis. [58,59]

Taken as a whole, these data suggest that there is a complex interaction between the metabolic activity of the VAT and the local immune cell fraction in SG animals. However, given both the metabolic and immune gene profiling outlined above, it is unclear if local adipocyte function is driving immune changes or if there is immunologic remodeling leading to increase glucose sequestration and utilization. As outlined by Solinas and colleagues, adipocyte processes, like DNL, are capable of driving both local and global metabolic and immune function via PPAR signaling and effecting GLP-1 secretion, hepatic lipogenesis, insulin sensitivity, and the host gut barrier and immunity. [60] [43,61] Alternatively, it is plausible that local immune remodeling is driving metabolism. Immune cell trafficking, activation, differentiation, and retention are all metabolically demanding. When activated, T cells change from a quiescent state reliant on basal oxidative phosphorylation, to a highly anabolic state with increased mitochondrial production as well as increased contributions of the pentose phosphate pathway, glycolysis, and glutaminolysis to energy production. [62,63] This increased demand for reducing equivalents may alter the glucose uptake of the VAT depot and drive the RER in the direction of CO_2_ production (i.e. above 1.0). Similarly, macrophage, T-cell, and B cell polarization and production of tolerance within dendritic cells can have profound effects upon metabolism. [64–66] [67] Finally, we have previously shown in rats that SG induces reductions in jejunal expression of IL-17, IL-23, and Interferon gamma, which are correlated with weight loss and systemic insulin levels in SG animals. [68]

Thus, it is plausible that VAT-specific changes underlie the SG-related improvements in glucose handling by increasing metabolically demanding local processes, which in turn affect global metabolism. In fact, studies of caloric restriction have shown that feeding can induce adipose tissue DNL, increase RER above 1.0, and lead to innate immune modulation. [26,69]

## 5. Limitations

Our study was completed in C57BL/6J mice fed normal rodent chow. Li et al., showed that SG animals in a DIO model, decreased the average RER toward 0.7. [70] These differences can be partially explained the available dietary resources in the experiment and may indicate that SG mice, when exposed to excess fat calories, adaptively increase beta oxidation. Further, in their experiment, calorimetry was performed at 7 weeks post-operatively and thus, it is possible that they missed the weight-loss independent effects that occur early in the post-surgical time course and that our finding of increased RER may become extinct as physiologic adaptation to post-surgical changes occur. The latter is in keeping with the clinical reality that the main physiologic driver of diabetes remission occurs within days of surgery and that further improvements may occur with weight loss. [1,2,5,71] Finally, given the nature of our model, we were not able to explore the weight-dependent effects of SG.

## 6. CONCLUSION

SG induces immediate and durable, weight-loss independent increases in glucose utilization heralded by enhanced metabolic activity of and local immunologic remodeling of VAT. Future studies should be aimed at better understanding the interplay of host immunity and adipocyte biology to post-SG physiology.

## ABBREVIATIONS

SG: Sleeve Gastrectomy
POD: post-operative Day
GLP-1: glucagon like peptide 1
OGTT: oral glucose tolerance test
ITT: insulin tolerance test
PET/CT: positron emission tomography-computed tomography
CLAMS: comprehensive lab animal monitoring systems
RER: respiratory exchange ratio
EE: energy expenditure
DNL: de novo lipogenesis
VAT: visceral adipose tissue
T2D: Type 2 Diabetes
FXR: farnesoid X Receptor
PPARγ: peroxisome proliferator-activated receptor gamma
FOXO1: forkhead box protein O1
SMRT: silencing mediator of retinoid and thyroid hormone receptors
NCOR: nuclear receptor co-repressor
MECOM: MDS1 and EVI1 Complex Locus
SUZ12: Polycomb Repressive Complex 2 Subunit
E2A: E2A immunoglobulin enhancer-binding factor E12/E47
MTF2: Metal Response Element Binding Transcription Factor 2
GATA3: gata binding protein 3

