## Appendices for "Sleeve Gastrectomy enhances glucose utilization and remodels adipose tissue independent of weight loss"

**APPENDICIES**

| **Table A1.** ANCOVA analysis from POD 7 to 14 (lean mass covariate) | | | |
| --- | --- | --- | --- |
|  | Full Day | Light | Dark |
| Respiratory Exchange Ratio | **<0.001** | **<0.001** | **0.0053** |

| **Table A2.** GLM analysis from POD 7 to 14 (lean mass covariate) | | | | | | | | | |
| --- | --- | --- | --- | --- | --- | --- | --- | --- | --- |
|  | Full Day | | | Light | | | Dark | | |
|  | Mass | Group | Interaction | Mass | Group | Interaction | Mass | Group | Interaction |
| Oxygen Consumption (ml/hr) | 0.0179 | 0.0315 |  | 0.0244 | 0.1419 |  | 0.0181 | 0.0096 |  |
| Carbon Dioxide Production (ml/hr) | 0.0491 | 0.3969 |  | 0.0494 | 0.7671 |  | 0.0637 | 0.2345 |  |
| Energy Expenditure (kcal/hr) | 0.0255 | 0.0473 |  | 0.0445 | 0.1305 |  | 0.0218 | 0.0251 |  |
| Hourly Food Consumed (g) | 0.0335 | 0.8854 |  | 0.0365 | 0.8639 |  | 0.0316 | 0.9033 |  |
| Total Food Consumed (g) | 0.3561 | 0.8790 |  | 0.3313 | 0.8466 |  | 0.3797 | 0.9075 |  |

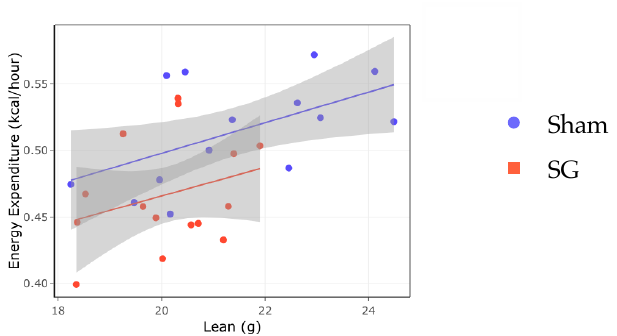

**Figure A3:** In the first post-operative week, SG mice have lower EE per gram of lean mass; ANCOVA with lean mass covariate; p = 0.026 for a 24-hour period; shadowing Std Error).

| **Table A4.** ANCOVA analysis from POD 27 to 34 (lean mass covariate for EE) | | | |
| --- | --- | --- | --- |
|  | Full Day (p-value) | Light (p-value) | Dark (p-value) |
| Energy Expenditure (kcal/hr) | 0.584 | 0.969 | 0.326 |
| Respiratory Exchange Ratio | **0.019** | **0.033** | **0.039** |

| **Table A5.**18-FDG SUV quantification at 90 minutes via well counter (SG n=4, Sham n=5) | | | | |
| --- | --- | --- | --- | --- |
|  | Sham | SG | p | |
| Whole Blood | 0.38±0.19 | 0.44±0.22 | | 0.65 |
| Stomach | 3.19±1.29 | 1.68±0.87 | | 0.06 |
| Duodenum | 3.63±1.63 | 1.93±0.72 | | 0.07 |
| Proximal Small Bowel | 1.61±1.64 | 1.40±0.56 | | 0.79 |
| Mid Small Bowel | 0.94±0.64 | 1.75±1.17 | | 0.36 |
| Distal Small Bowel | 0.76±0.42 | 2.22±2.21 | | 0.18 |
| Cecum | 0.99±0.43 | 1.55±0.64 | | 0.14 |
| Colon | 0.57±0.30 | 1.29±0.78 | | 0.20 |
| **Cecal Stool** | **0.04±0.03** | **0.28±0.08** | | **0.007** |
| Liver | 0.50±0.20 | 0.70±0.26 | | 0.23 |
| Pancreas | 0.32±0.18 | 0.47±0.17 | | 0.26 |
| Kidney | 1.22±0.70 | 1.85±0.92 | | 0.28 |
| Spleen | 0.62±0.36 | 1.19±0.57 | | 0.11 |
| Heart | 6.77±3.14 | 9.92±3.70 | | 0.21 |
| **Lung** | **1.09±0.38** | **1.83±0.48** | | **0.04** |
| Thigh Muscle | 0.10±0.04 | 0.13±0.04 | | 0.30 |
| Testes | 0.34±0.17 | 0.58±0.27 | | 0.15 |
| Whole Brain | 0.66±0.33 | 1.02±0.42 | | 0.31 |
| Femur | 1.05±0.47 | 2.01±0.79 | | 0.15 |
| Adrenals | 1.20±0.78 | 2.26±0.89 | | 0.19 |
| Bladder | 6.57±2.13 | 12.17±11.5 | | 0.37 |
